# CRISPR-PTM and CRISPR-VEIS: Multiplexed platforms for quantitative functional analysis of endogenous phosphosites

**DOI:** 10.64898/2026.05.07.723463

**Authors:** Simon Willaume, Jan Benada, Karen Akopyan, Valdemaras Petrosius, Jayashree Vijay Thatte, Thomas C.R. Miller, Arne Lindqvist, Claus S. Sørensen

**Affiliations:** Biotech Research and Innovation Center, University of Copenhagen; Department of Cell and Molecular Biology, Karolinska Institutet, Biomedicum, Stockholm, Sweden; DNRF Center for Chromosome Stability, Department of Cellular and Molecular Medicine, University of Copenhagen

## Abstract

Connecting protein post-translational modifications (PTMs) to phenotypic outcomes is a central challenge. Although phosphoproteomics has richly catalogued specific sites, reliable methods to measure the endogenous effects of individual phosphosites on cellular fitness and signaling are still lacking. Here, we introduce CRISPR-PTM and CRISPR-VEIS as complementary platforms for quantitative, endogenous phosphosite interrogation at both individual and clustered phosphorylation events. CRISPR-PTM is a multiplexed knock-in framework generating defined phosphosite variants with internal allelic markers, enabling precise relative fitness effects in pooled populations. CRISPR-VEIS (Visualisation of Edits In Situ) is an in situ mRNA-genotyping approach that directly links endogenous allelic edits to single-cell phenotypes, addressing needs for subclonal isolation or exogenous reporters. We applied these methods to the WEE1–CDK1 regulatory pathway, where canonical CDK1-Y15 phosphorylation alone cannot explain WEE1 loss or inhibition phenotypes. CRISPR-PTM systematically quantified fitness consequences of CDK1 phosphosite variants and identified Y19 as a previously unrecognized WEE1-dependent inhibitory site. Single non-phosphorylatable substitutions at Y15 or Y19 had minimal impact, but combined CDK1-Y15F/Y19F editing caused pronounced fitness defects, phenocopying WEE1 inactivation and showing epistasis to WEE1 inhibitors. CRISPR-VEIS further demonstrated that acute endogenous editing of both sites correlated with elevated CDK activity at the single-cell level. Together, CRISPR-PTM and CRISPR-VEIS provide broadly applicable approaches for quantitative analysis of PTM function, enabling direct linkage of endogenous phosphosite variation to cellular fitness and signaling phenotypes.

## Introduction

PTMs are central regulators of protein activity and signaling, yet directly and precisely connecting PTM events to cellular phenotypes remains difficult. Large-scale phosphoproteomics has catalogued tens of thousands of sites^1^, but for most phosphosites we lack quantitative, endogenous measurements of how PTMs affect key aspects like cellular fitness, signaling dynamics, or drug responses^2^. This problem is compounded when PTMs occur in clusters or when perturbations produce subtle or context-dependent effects that are masked by population averaging.

Existing experimental strategies each have important limitations. Overexpression of mutant constructs distorts protein stoichiometry and localization; generation and characterization of clonal genome-edited lines is slow and can be misleading for subtle or essential impact PTMs; pooled CRISPR screens targeting enzymes provide broad discovery power but generally lack single-site, quantitative resolution and cannot directly link endogenous phosphosite alleles to single-cell phenotypes^3^. Recent massive base editing approaches have markedly expanded functional phosphosite interrogation. However, they display edit-context biases and are optimized for discovery lacking hypothesis testing opportunities, thus, they cannot deliver precise, multiplexed interrogation of adjacent sites ^4–7^. Finally, there is a lack of methods that directly combine in situ detection of knock-in edited cells with cell-based phenotype markers, which hampers deconvolution of phosphosite functions.

To address these gaps, we developed two complementary CRISPR-based platforms: CRISPR-PTM and CRISPR-VEIS. CRISPR-PTM is a multiplexed knock-in framework that introduces defined phosphosite variants -non-phosphorylatable, phospho-mimetic and an internal synonymous wild-type control -in parallel at endogenous loci, including editing multiple sites in vicinity. By tracking changes in allelic ratios over time by deep sequencing and fitting growth models, we obtain sensitive, quantitative estimates of variant-specific fitness in pooled populations. CRISPR-VEIS (Visualization of Edits In Situ) adapts allele-specific probes and rolling-circle amplification to detect edited mRNA in fixed cells, enabling direct in situ linkage between endogenous alleles and phenotypic markers at single-cell resolution without clonal isolation. Moreover, it discriminates CRISPR knock-in alleles with single-nucleotide discrimination. To unequivocally demonstrate CRISPR-PTM and CRISPR-VEIS potential, we deployed them to unravel a long-standing mechanistic enigma within the indispensable and clinically relevant CDK1 regulatory pathway. CDK1 signaling governs the cell cycle being tightly controlled by the kinases WEE1 and PKMYT1, which phosphorylate tyrosine 15 (Y15) and threonine 14 (T14), respectively^8–13^. Small molecule inhibitors of WEE1 and PKMYT1, such as adavosertib^14^, zedoresertib (Debio 0123)^15^, and lunresertib^16,17^, have shown anticancer effects alone or in combination by inducing supraphysiological CDK activity that leads to cell death^17–21^. Although both PKMYT1 and WEE1^22^ kinases are essential, individual non-phosphorylatable point mutations of CDK1 (T14A or Y15F) display little or no observable phenotypes in standard functional assays^23–25^.

We applied CRISPR-PTM and CRISPR-VEIS to dissect CDK1 regulatory phosphorylation and its connection to WEE1 inhibition. CRISPR-PTM delivered sensitive, multiplexed fitness readouts for single and combined CDK1 phosphosite variants, while CRISPR-VEIS linked endogenous edits to CDK activity at single-cell resolution. Using these complementary approaches, we identified a cooperative inhibitory mechanism based on WEE1 mediated dual-site phosphorylation of CDK1 that together underpins cell fitness and determines responses to WEE1 inhibitors. Overall, this work provides a powerful experimental framework to functionally decode signaling events to downstream phenotypes

## Results

### CRISPR-PTM enables multiplexed, quantitative fitness analysis of CDK1 phosphorylation-site variants

To address the lack of quantitative phenotype assays for single and clustered PTM knock-in, we developed CRISPR-PTM. This is a multiplexed ssODN knock-in framework with an internal synonymous WT’ control that allows direct, allele-frequency based estimation of relative fitness in pooled populations. With focus on phosphosites, we simultaneously introduced multiple defined phosphosite variants within the same culture system, including non-phosphorylatable (e.g., alanine/phenylalanine substitutions), phospho-mimetic (e.g., aspartic acid/glutamic acid substitutions), and co-edited allelic synonymous control (WT’) (Fig. 1a). This new approach is based on the simple CRISPR-Select assay, where only a single variant of interest is tracked against one synonymous variant^26^. We first set out to validate its precision, reproducibility, and modeling workflow below. As a proof-of-concept case we focused on CDK1, a key cell cycle regulator whose activity is controlled by two adjacent N-terminal phosphorylation sites, T14 and Y15. Given that the functional logic of the CDK1 pathway remained unclear, we set out to quantify the fitness effects (Fig. S1a) using CRISPR-PTM. To test the multi-site knockin assay on CDK1, mutation effects were first assessed with focus on a temporally extended format in non-transformed breast epithelial cells (MCF10A) engineered to be p53-deficient^27^. We edited CDK1-T14 and CDK1-Y15 to generate desired variants (Fig. 1b,c), and quantified mutant/WT’ allele ratios by allelic deep sequencing at intervals for a total of 18 days (Fig. S1b). CDK1-T14A modestly enhanced proliferation^23^, whereas CDK1-Y15F had no detectable fitness impact despite the well-established cytotoxicity of WEE1 inhibition or loss^14,18,22^. Phospho-mimetic CDK1-T14D and CDK1-Y15E strongly impaired fitness, consistent with inhibitory phosphorylation of CDK1 by PKMYT1 and WEE1^28–30^ (Fig. 1b,c). Our CRISPR-PTM measurements were highly reproducible across biological replicates: replicate Mut:WT′ allele-frequency profiles correlated with Pearson r = 0.95– 0.99 (n = 3–6), and estimated fitness values showed a typical coefficient of variation <10% when experiments met the sequencing and editing thresholds described in Methods. To assess the temporal appearance of phenotypes, we introduced the same mutations in a short-term assay and included the combined T14A/Y15F substitution (CDK1-AF). With this approach, we tested multi-site editing of adjacent sites and explored a positive control for synthetic lethality with an expected rapid appearing phenotype (Fig. 1d,e)^31,32^. CDK1-AF was well edited and displayed lethality, markedly reducing fitness within 3–4 days. This matched prior reports of synthetic lethality between WEE1 and PKMYT1 inhibition^18,19^. Short-term measurements mirrored long-term trends: CDK1-Y15F lacked detectable effect, CDK1-T14A conferred a mild fitness advantage, and phospho-mimetic variants produced strong phenotypes (Fig. 1d and 1e).

**Figure 1:**
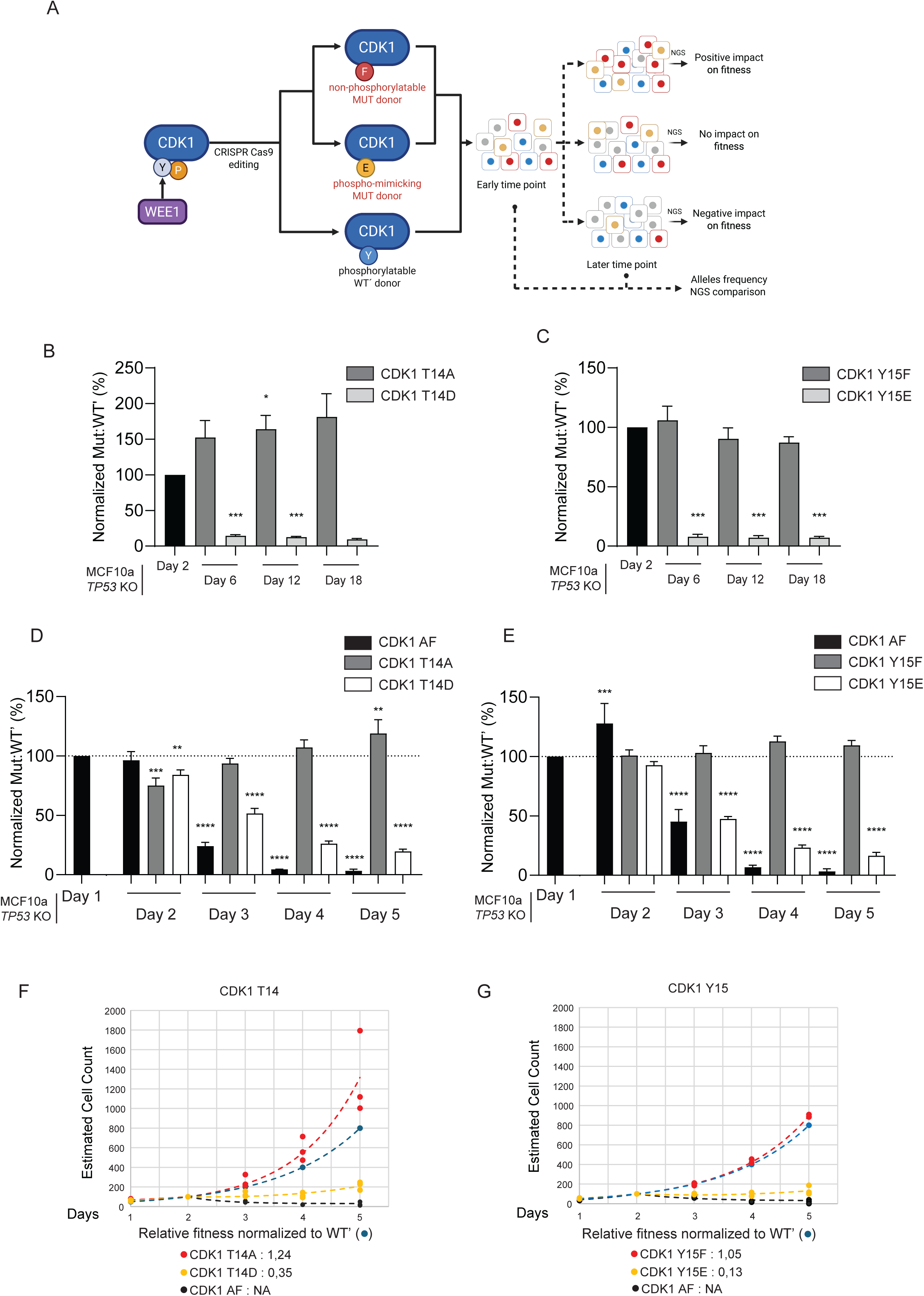
CRISPR-PTM quantifies fitness effects of CDK1 G-loop phosphorylation sites. (A) Strategy for quantitative phosphosite editing and fitness determination with CRISPR-PTM. Endogenous phosphosites are edited to non-phosphorylatable (A/F) or phospho-mimetic (D/E) residues and co-edited with a synonymous, phosphorylatable internal control (WT′). Relative fitness is inferred from changes in Mut/WT′ allele ratios between early and late time points. As examples, the CDK1-Y15 site, targeted by the WEE1 kinase, is replaced with phenylalanine and glutamic acid. (B) CDK1-T14 phosphosite variants show opposite fitness effects. CRISPR-PTM analysis of CDK1-T14A and T14D in iCas9-MCF10A TP53KO cells. Mut/WT′ ratios were measured at days 2, 6, 12, and 18 and normalized to day 2. Mean ± SEM; n = 3. (C) CDK1-Y15 phosphosite variants display differential fitness effects. CRISPR-PTM analysis of CDK1-Y15F and Y15E variants was performed and analyzed as in (B). Mean ± SEM; n = 3. (D) CRISPR-PTM quantifies short-term fitness impact of CDK1-T14 mutations based on short time-resolved analysis (days 1–5) of CDK1-T14A and T14D. CDK1-T14A/Y15F (AF) serves as a positive control. Data are normalized to day 1 (dotted line). Mean ± SEM; n = 5. (E) Short-term fitness impact of CDK1-Y15 mutations measured as in (D) for CDK1-Y15F and Y15E variants. Mean ± SEM; n = 5. (F, G) Modeling of data from CRISPR-PTM. Experimental read counts for wild-type and mutant alleles were converted to estimated cell numbers, assuming 100 cells at day 2. Observed data points (solid circles) are shown alongside fitted curves (dashed lines) from two-phase growth models (see Materials and Methods). Relative fitness of mutant compared to wild-type alleles is shown for day 2-5.

We next sought to extract a quantitative measure of the fitness change after CRISPR-PTM. We estimated population size from sequencing reads and noted that the data can be divided into two patterns, before and after day 2 (Fig. 1f and 1g). We therefore fitted the data to a two-phase growth model^33^, in which the first days are fitted to an exponential growth pattern, and the following days are fitted to a model in which exponential growth is combined with decay. The combination of growth and decay components indicate that the model can fit a population in which both proliferation and ceased proliferation are present. Indeed, the data fittings would be consistent with phospho-mimetic CDK1 variants showing reduced fitness both due to extended cell cycle duration and to a subpopulation that does not proliferate after day 2. In contrast, the data would be consistent with a model in which reduced cell cycle duration is a primary cause of increased fitness of non-phosphorylatable CDK1 variants, particularly for CDK1-T14A.

Next, because p53 has been linked to the WEE1–CDK1 pathway^14^, we repeated fitness assays in the parental *TP53*-proficient breast epithelial cell line. CDK1-T14A and CDK1-Y15F variants behaved similarly in wild-type and *TP53*-null MCF10A cells (Fig. S1c). This indicated limited p53 involvement in phenotypes, thus, we conducted ensuing experiments in *TP53*-deficient cells. To further probe potentially concealed phenotypes for non-phosphorylatable CDK1-Y15F, we treated CRISPR knock-in populations with the PKMYT1 inhibitor lunresertib. CDK1-Y15F cells were strongly sensitized to lunresertib (Fig. S1d–f), which was fitting with a conditional phenotype unmasked by dual inhibition of CDK1 inhibitory kinases. Together, these data establish CRISPR-PTM as a precise and sensitive platform for functional PTM analysis providing multisite results. Moreover, the data reinforced the longstanding paradox that CDK1-Y15F alone fails to recapitulate the essential role of WEE1.

### Phosphosite editing reveals incomplete epistasis between CDK1-Y15F and WEE1 inhibition

With the CRISPR-PTM system established, we next quantified genotype–drug interactions between CDK1 phosphorylation-site edits and WEE1 inhibition (Fig. 2a). If phosphorylation of CDK1-Y15 was the sole functionally relevant output of WEE1, a non-phosphorylatable CDK1-Y15F variant would be expected to phenocopy WEE1 inhibition and to be epistatic to WEE1 inhibitors. We first applied adavosertib, which has undergone multiple clinical trials^14,34,35^. Editing CDK1-T14A followed by treatment with a tolerable dose of adavosertib (250 nM, Fig. S1d) led to marked depletion of CDK1-T14A knock-in cells (Fig. S1d,e). This was in agreement with the synthetic lethality between WEE1 and PKMYT1 inhibition. Importantly, CDK1-Y15F cells that mimic loss of WEE1 activity at Y15 remained moderately sensitive to adavosertib (Fig. 2b). This lack of epistasis suggested either additional CDK1 phosphorylation sites under WEE1 control, or that non-CDK1 targets were contributing to WEE1 inhibitor responses. Because WEE1 also phosphorylates CDK2-Y15^36^ (Fig. S1a), we examined whether CDK2 activity might mask the phenotype of CDK1-Y15F. First, we conducted knock-in editing of the CDK2 T14/Y15 sites, however, this showed only a minor, non-significant fitness effect for the phospho-deficient CDK2-T14A/Y15F variant (Fig. 2c). Next, to shift cell-cycle dependence toward CDK1, we generated MCF10A *TP53*KO cells that also lack *CDK2*^28,29,37^ (Fig. S2a). In drug response assays, these cells retained synthetic lethality to combined WEE1 and PKMYT1 inhibition, although the effects were slightly attenuated (Fig. S2b,c and Fig. S1d). We then assessed CDK1 phosphorylation-site edits in the CDK2-deficient, more CDK1-reliant background. In this setting, phosphosite phenotypes were augmented and CDK1-Y15F yielded a moderate fitness effect (Fig. 2d). Notably, WEE1 inhibition still did not display epistasis with the CDK1-Y15F knock-in in (Fig. 2e). Thus, CDK2 does not fully account for the persistent sensitivity of CDK1-Y15F cells to WEE1 inhibition. This incomplete epistasis further indicated that Y15 phosphorylation cannot fully account for the essential role of WEE1.

**Figure 2:**
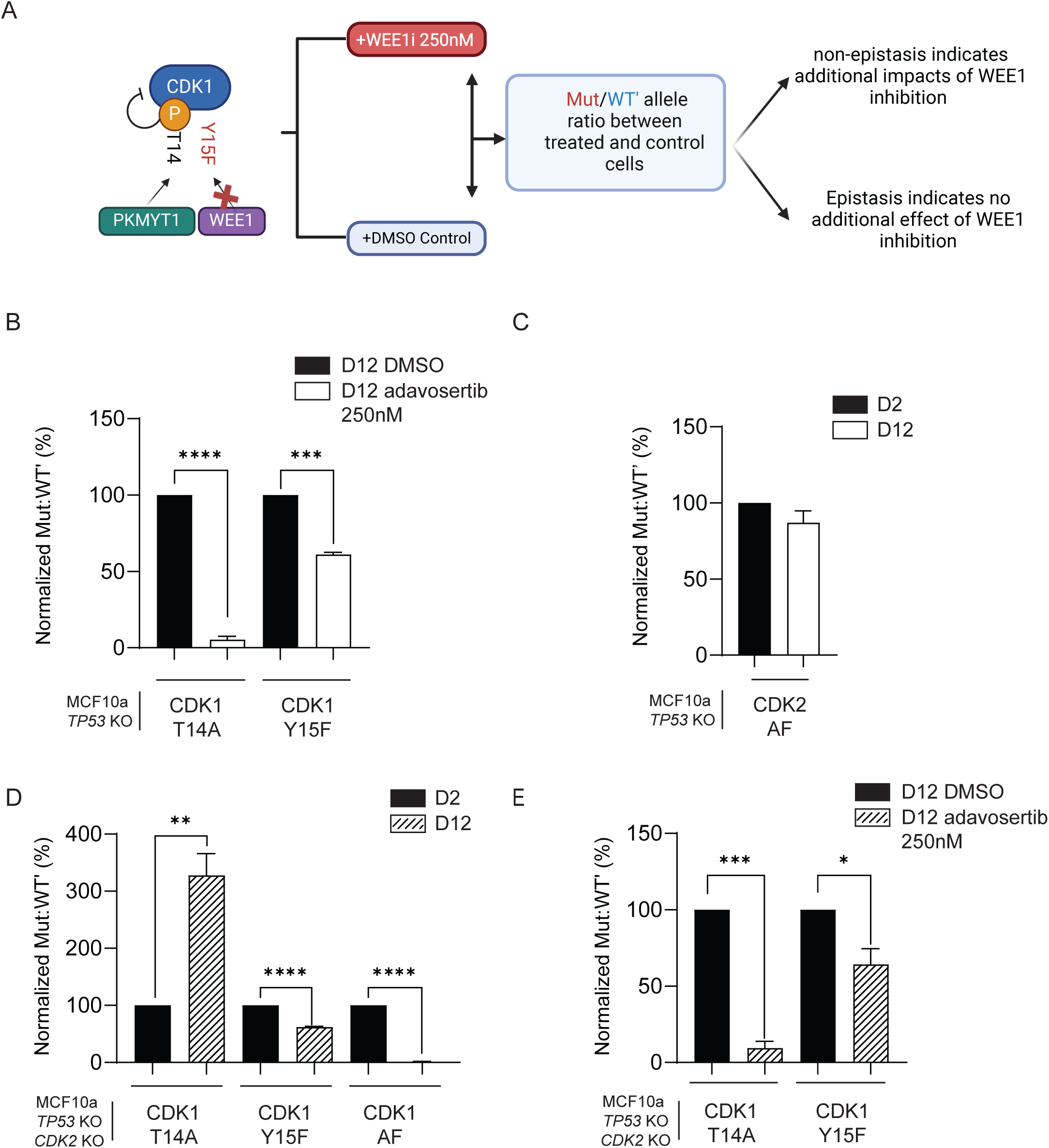
Epistasis analysis between WEE1 inhibition and CDK1-Y15 phosphosite variants. (A) CRISPR knock-in epistasis analysis framework for kinase inhibition. A non-phosphorylatable mutation at a target site for a specific kinase is combined with chemical inhibition of the kinase. The WEE1-CDK1 pathway is shown as example, lack of additional fitness effects upon inhibition indicates epistasis. (B) CDK1-Y15F mutation is not epistatic with adavosertib WEE1 inhibition. CRISPR knock-in analysis of CDK1-T14A and Y15F variants treated with DMSO or adavosertib (250 nM). Mut:WT′ ratios were determined on day 12 treated with either DMSO as control or adavosertib, and normalized to day 12 DMSO value. Mean ± SEM; n = 4. (C) Non-phosphorylatable CDK2-T14A/Y15F has limited fitness impact. CRISPR knock-in analysis of CDK2-T14A/Y15F variant over time, normalized to day 2 value. Mean ± SEM; n = 4. (D) CDK1 phosphosite phenotypes are exacerbated by CDK2 ablation. CRISPR knock-in analysis of CDK1-T14A, Y15F, and AF variants in iCas9-MCF10A, *TP53*KO, *CDK2*-KO cells. Data are normalized to day 2 value. Mean ± SEM; n = 6. (E) CDK1-Y15F mutation and WEE1 inhibition remain non-epistatic in CDK2 deficient cells. Experiments were conducted as in (B) with iCas9-MCF10A, TP53KO, CDK2-KO cells. Mean ± SEM; n = 4. P values are from Paired t-test with * = 0.0332, ** = 0.0021, *** = 0.0002, and **** = 0.0001.

### CDK1-Y19/Y15 are WEE1-regulated inhibitory sites jointly required to ensure fitness

Given that CDK1-Y15F edited cells displayed mild phenotypes and remained sensitive to WEE1 inhibition, we applied an unbiased, short-term phosphoproteomic screen to systematically identify primary WEE1-dependent phosphorylation sites. We used a controlled 1.5 h treatment with DMSO, adavosertib or lunresertib to prioritize kinase substrates while minimizing secondary, cell-cycle–driven changes for follow-up validation (Fig. S3a). This analysis confirmed the known WEE1-dependent phosphorylation of CDK1-Y15 and identified CDK1-Y19 as a previously uncharacterized site. The Y19 site is proximal to the G-loop and highly conserved across species (Fig. 3a). While reported as phosphorylated in the PhosphositePlus^1^ database, CDK1-Y19 had not previously been linked to WEE1 or functionally investigated. To further our analysis, we generated phospho-specific antibodies targeting this residue. CDK1-Y19 phosphorylation was detectable in MCF10A cells after immunoprecipitation (Fig. 3b), as well as endogenously in U2OS cells (Fig. S3b-d). Importantly, WEE1 inhibition or CDK1 depletion led to a decrease in CDK1-Y19 phosphorylation (Fig. 3b, Fig. S3c,d). To assess its functional relevance, we applied knock-in editing to generate phospho-mimetic and non-phosphorylatable variants. Phospho-mimetic mutations (CDK1-Y15E, Y19E, Y15E/Y19E) revealed that Y19E alone moderately reduced cell fitness, whereas Y15E and the double Y15E/Y19E mutations had stronger effects (Fig. 3c). Moreover, non-phosphorylatable CDK1-Y19F alone had little impact (Fig. 3d). Importantly, fitness phenotypes were observed in the CDK1-Y15F/Y19F mutant indicating a phenotype akin to WEE1 deficiency (Fig. 3e). We then explored the functional responses of phosphosite mutants to PKMYT1 or WEE1 inhibition. CDK1-Y19F cells were sensitive to PKMYT1 inhibition (Fig. s3e). In addition, combining CDK1-Y19F and T14A significantly impaired fitness relative to CDK1-T14A alone in *CDK2*-wt (comparing Fig. S3f and Fig. 1b). A fitness impact was also observed in *CDK2*-KO cells (comparing Fig. S3f with Fig. 2d). Based on this, we reasoned that CDK1-Y19 could be important for cellular WEE1 inhibitor responses. When conducting functional analysis, we observed that CDK1-Y19F displayed moderate sensitivity to WEE1 inhibition both in the presence and absence of CDK2 (Fig. 3f). This lack of epistasis indicated a WEE1 impact beyond CDK1-Y19. We then tested the dual CDK1-Y15F/Y19F mutation in CDK2-deficient cells that are CDK1 reliant. Here, the combined mutant was fully epistatic to WEE1 inhibition by adavosertib and zedoresertib^26^ (Fig. 3g and 3h). Altogether, these results suggest that WEE1 controls CDK1 via residues Y15 and Y19, and that CDK2 contributes to a subset of the fitness defects observed upon WEE1 inhibition.

**Figure 3:**
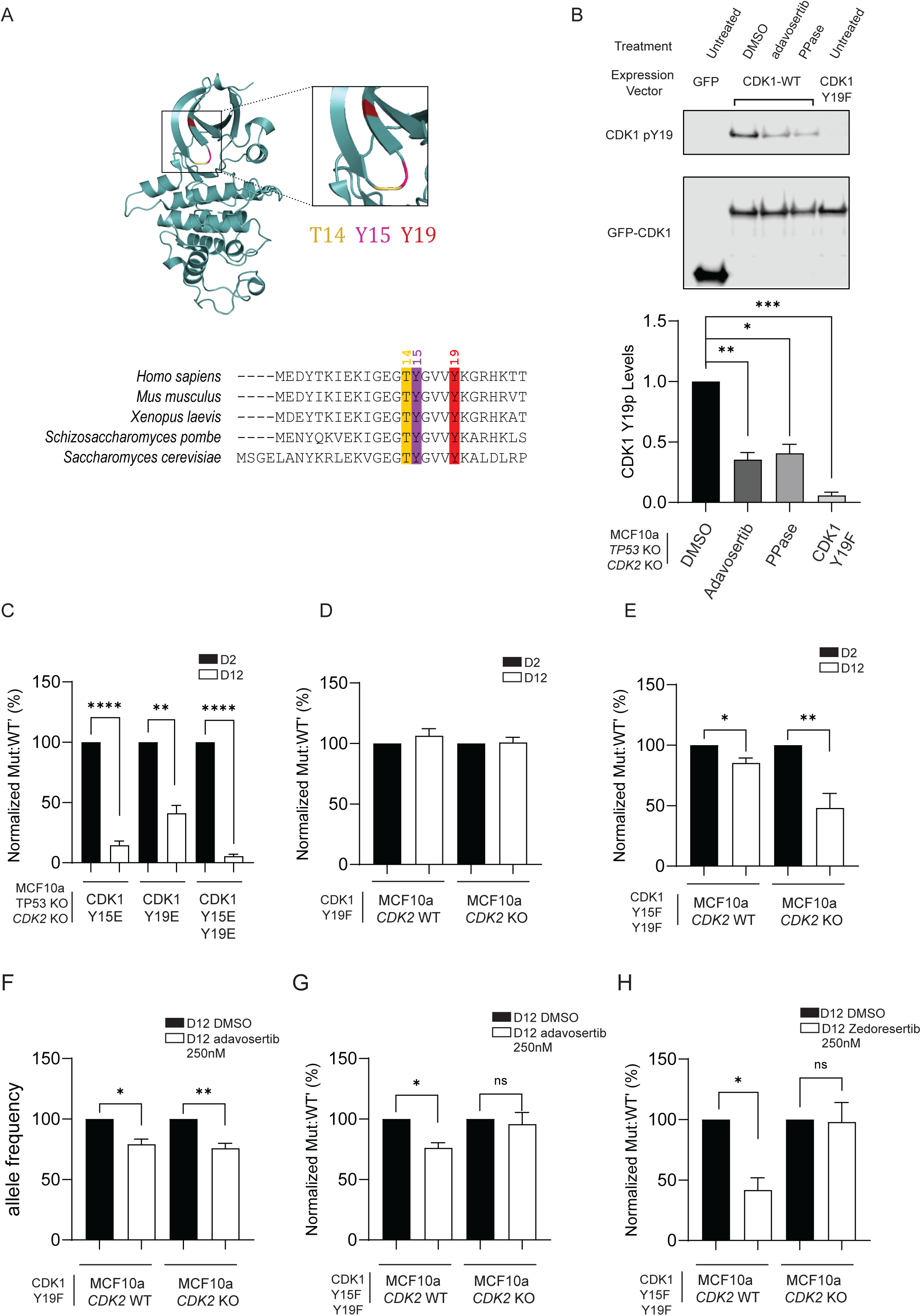
WEE1 mediated phosphorylation of CDK1-Y19 promotes fitness. (A) CDK1-Y19 is a conserved G-loop residue. Structural representation of the CDK1 G-loop highlighting T14, Y15, and Y19 (top), and sequence conservation across species (bottom). The structure is based on https://doi.org/10.2210/pdb6gu2/pdb protein structure, visualized in The PyMOL Molecular Graphics System, version 2.5.1 -https://.pymol.org/. The comparison of protein sequences was done using Clustal Omega Multiple Sequence Alignment (MSA) -https://.ebi.ac.uk/jdispatcher/msa/clustalo. (B) CDK1-Y19 phosphorylation is WEE1-dependent. IP-immunoblot experiments were done in iCas9-MCF10A, *TP53*KO, *CDK2* KO, inducible for CDK1-WT-GFP or CDK1-Y19F-GFP or just GFP as a control. Cells were treated with WEE1i (adavosertib; 1 μM, 4 h, lane 3). Phosphorylation was impacted by treating GFP-IP proteins with lambda phosphatase for 30 min. Image shows a representative immunoblot result after GFP-IP. Lower part data are normalized to DMSO conditions and depict means ± S.E.M of n = 3 independent biological replicates. (C) Phospho-mimetic CDK1-Y19 substitutions reduce cell fitness. CRISPR knock-in analysis of CDK1-Y15E, Y19E, and Y15E/Y19E variants in iCas9-MCF10A cells *TP53*KO *CDK2* KO. Data are normalized to day 2 value. Mean ± SEM; n = 3. (D) Individual phosphosite editing of non-phosphorylatable CDK1-Y19F does not reduce cell fitness. CRISPR knock-in analysis was carried out in iCas9-MCF10A, *TP53*KO cells that were either *CDK2* WT or *CDK2* KO. Data are normalized to day 2 value. Mean ± SEM; n = 5. (E) Marked fitness impact of combined non-phosphorylatable CDK1-Y15/Y19 mutant. CRISPR knock-in analysis of CDK1-Y15F/Y19F variants was carried out in iCas9-MCF10A, *TP53*KO cells that were either *CDK2* WT or *CDK2* KO. Data are normalized to day 2 value. Mean ± SEM; n = 5. (F) Non-epistasis between CDK1-Y19F and WEE1 inhibition. CRISPR knock-in analysis of Y19F variants treated with DMSO or adavosertib (250 nM). Data are normalized to day 12 DMSO value. Means ± S.E.M; n = 4 (**G–H)** Dual non-phosphorylatable CDK1-Y15/Y19 mutants are epistatic to WEE1 inhibition in CDK2 deficient cells. CRISPR knock-in analysis of Y19F or Y15F/Y19F variants treated with DMSO, adavosertib or zedoresertib (250 nM). Data are normalized to day 12 DMSO value. Mean ± SEM; n = 3–4. P values are from Paired t-test with * = 0.0332, ** = 0.0021, *** = 0.0002, and **** = 0.0001.

### CRISPR-VEIS links acute endogenous CDK1-Y15/Y19 edits to elevated CDK activity at single-cell resolution

To move beyond population averages and directly link endogenous allelic edits to immediate cellular phenotypes, we complemented CRISPR-PTM with CRISPR-VEIS. This is an allele-specific mRNA genotyping approach that detects knock-in edits with a fluorescent signal allowing single-cell phenotype analysis. Below we describe probe design, specificity controls, and how CRISPR-VEIS recapitulates the population-level findings at single-cell resolution with a focus on CDK1-Y19. To avoid potentially artefact-prone and tedious subcloning experiments, we aimed instead to detect edited cells directly in situ while preserving the design principle of comparing simultaneously generated mutations of interest with synonymous controls in the same culture system. We reasoned that allele-specific detection of edited mRNA would provide the most accurate readout, given challenges in identifying a single genomic edit in situ at the chromosome level. To this end, we developed CRISPR-VEIS which is based on padlock probe detection coupled with rolling circle amplification (RCA) for in situ RNA detection^38^. This allows simultaneous microscopy analysis of cell-based markers for genotype and phenotypes (Fig. 4a). As proof of concept, padlock-RCA robustly detected edited CDK1 transcripts in MCF10A cells, whereas only a rare signal was observed in non-edited controls (Fig. 4b; Fig. S4a). Moreover, editing frequencies were rather similar when compared between padlock-RCA and NGS-based quantification (Fig. S4b), confirming the accuracy of in situ genotyping. We then monitored CDK activity by quantifying phosphorylated FOXM1(T600), a canonical CDK target^39^. In situ analysis showed that CDK1-Y15F knock-in alone or combined with PKMYT1 inhibition increased CDK activity relative to synonymous controls (Fig. 4c). Importantly, CDK1-Y19F knock-in led to an increase in CDK activity as seen for CDK1-Y15F (Fig. 4d). Together, this showed that CRISPR-VEIS can enable direct, quantitative genotype–phenotype mapping in situ and demonstrates that WEE1-mediated phosphorylation of CDK1-Y19, like CDK1-Y15, contributes to the control of CDK catalytic activity.

**Figure 4:**
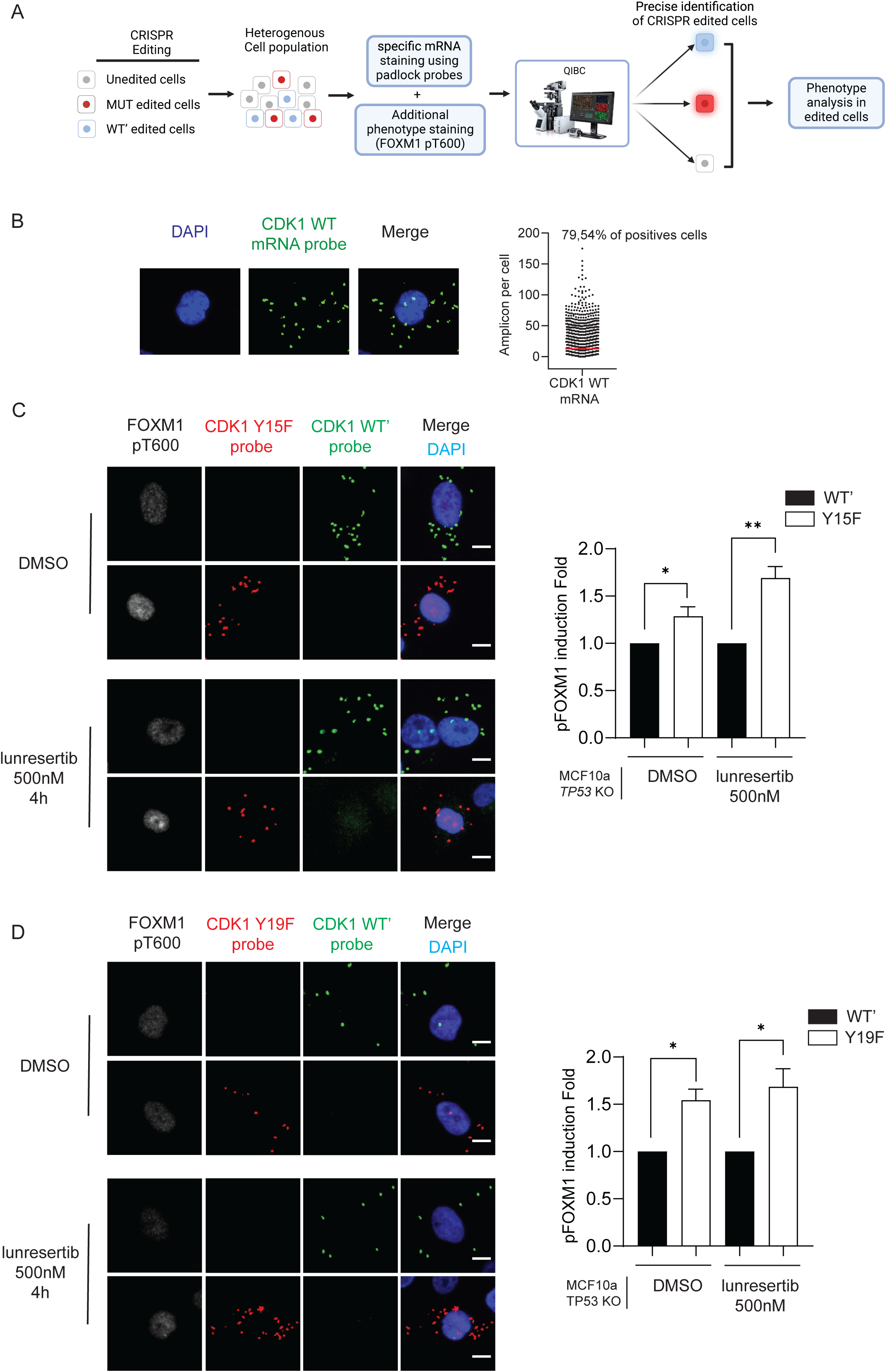
CRISPR-VEIS links CDK1 phosphosite genotypes to single-cell CDK activity. (A) CRISPR-VEIS workflow for in situ genotype–phenotype coupling. Padlock probe-based RNA detection distinguishes mutant and WT′ alleles, enabling phenotype quantification in individually edited cells. Simultaneous microscopy phenotype analysis of cell-based markers is typically carried out using QIBC (Quantitative Image-Based Cytometry). (B) Single-cell detection of CDK1 transcripts by padlock probe coupled with rolling circle amplification (RCA). Left, Representative image of CDK1 WT mRNA RCA signal in MFC10A iCas9, *TP53*KO cells. Right, Quantification of the number of Amplicons per cell for CDK1 WT mRNA. Representative quantification of n = 3 individual biological replicates. (C) CDK1-Y15F enhances FOXM1 phosphorylation in situ. Left, Representative image of CDK1-Y15F (green) or WT’ (red) mRNA RCA signal in iCas9-MCF10A cells *TP53*KO cells treated with DMSO or lunresertib 500 nM for 4h. Cells were stained for FOXM1 pT600 (grey) and counterstained with DAPI (blue). Right, quantification of FOXM1 pT600 levels in cells population positive for FOXM1 pT600. Y15F data are normalized to the corresponding WT’. At least 150 edited cells were analyzed by experiment. n = 5 independent biological replicates. (D) Non-phosphorylatable CDK1-Y19F enhances FOXM1 phosphorylation in situ. Left, Representative image of CDK1-Y19F (green) or WT’ (red) mRNA RCA signal in MCF10A iCas9, *TP53*KO cells. Cells were treated and stained as in (Fig. 4c). Y19F data were normalized to the corresponding WT’. At least 150 edited cells were analyzed per replicate. n = 3 independent biological replicates. P values are from Paired t-test with * = 0.0332, ** = 0.0021, *** = 0.0002, and **** = 0.0001.

## Discussion

In this work, we introduce CRISPR-PTM and CRISPR-VEIS as complementary platforms for quantitative phosphosite interrogation. CRISPR-PTM leverages multiplexed knock-in and internal allelic controls to quantify the fitness effects of up to several nearby predefined phosphosite variants in pooled populations, while CRISPR-VEIS couples allele-specific in situ mRNA detection to phenotypic markers at single-cell resolution. Together, these methods provide a sensitive framework for connecting specific phosphosite changes to cellular outcomes in their native genomic context.

Applying these tools to the WEE1–CDK1 axis resolved a longstanding discrepancy between the essentiality of WEE1 and the unexpectedly mild phenotypes of non-phosphorylatable CDK1-Y15 mutants^23^. CRISPR-PTM revealed that WEE1 inhibition is not fully epistatic to CDK1-Y15F, and established CDK1-Y19 as an additional inhibitory site whose phosphorylation depends on WEE1. While single Y15F or Y19F substitutions had limited impact on cell fitness, combined CDK1-Y15F/Y19F editing strongly impaired proliferation and was epistatic to multiple WEE1 inhibitors. CRISPR-VEIS further demonstrated that edits at these sites acutely elevated CDK activity in individual cells. These findings support a dual-site inhibition model in which WEE1-mediated phosphorylation of CDK1 at Y15 and Y19 acts cooperatively to restrain CDK activity and maintain cell viability. Moreover, our research highlights a related conundrum for the PKMYT1-CDK1 pathway, where PKMYT1 is essential but CDK1-T14A even has a growth-promoting phenotype (Fig. 1b,d,f)^40^. Future studies will be needed to decode this, likely with the use of CRISPR-PTM and CRISPR-VEIS.

Beyond the WEE1–CDK1 pathway, CRISPR-PTM and CRISPR-VEIS are well-suited to dissect PTM function across diverse signaling networks. CRISPR-PTM can map fitness landscapes of clustered or combinatorial sites, functionally annotate disease-associated variants, and quantify how kinase inhibitors alter the contribution of specific sites to cell survival. CRISPR-VEIS provides spatially resolved genotype–phenotype mapping, enabling study of how PTM variants affect heterogeneous single-cell responses, such as pathway activation or drug tolerance. Both platforms are compatible with multiple phenotypic readouts and can be adapted beyond phosphorylation to other modifications (for example lysine acetylation) or defined proteolytic events. Their rapid, quantitative outputs make them suitable to map multiple-site interactions, annotate coding disease-related variants in their native context, explore essential variants and pathways, dissect site-specific drug mechanisms, and link edited genotypes to single-cell phenotypes. CRISPR-VEIS can also detect any defined endogenous edit (for example cancer-associated substitutions), not only PTMs.

These methods offer a faster, subcloning-free alternative to overexpression or clonal isolation, allowing physiologically relevant readouts at population and single-cell resolution. There is a subset of practical limitations linked to genome editing aspects and imaging setups. First, the multiplexing capacity of CRISPR-PTM is constrained by editing efficiency, repair biases, and available sequencing depth, which together limit the number of variants and loci that can be interrogated simultaneously. Second, CRISPR-VEIS currently requires high-content imaging and image analysis, which restricts throughput but is amenable to automation and parallelization. Ongoing advances in genome editing efficiency, barcoded probe libraries, and automated microscopy are expected to expand the scale and generalizability of CRISPR-PTM and CRISPR-VEIS.

## Abbreviations

CDK: Cyclin-Dependent Kinase
CDK1 AF: CDK1 T14A/Y15F
CDK1 DE: CDK1 T14D/Y15E
CRISPR: Clustered Regularly Interspaced Short Palindromic Repeats
EdU: 5-ethynyl-2’-deoxyuridine
FOXM1: Forkhead Box M1
PPase: Phosphatase
PKMYT1: Protein Kinase, Membrane Associated Tyrosine/Threonine 1
PTM: Post-Translational Modifications
QIBC: Quantitative Image-Based Cytometry
RCA: Rolling Circle Amplification
ssODN: Single-stranded oligodeoxyribonucleotide
VEIS: Visualization of Edit In Situ
WEE1: WEE1 G2 Checkpoint Kinase

## Acknowledgments

We thank the members of the CSS lab for insightful comments and fruitful discussion, Julien Duxin (BRIC, Copenhagen University) for critical feedback on the manuscript, and Sian Li (Brakebusch Group, BRIC, Copenhagen University) for the automated CRISPResso macro.

## Author contributions

Conceptualization: SW, JB, CSS

Methodology: SW, JB, KA, VP, JT, TM, AL, CSS

Investigation: SW, JB, KA, VP, JT, TM

Visualization: SW, JB, KA

Funding acquisition: CSS

Project administration: CSS

Supervision: CSS

Writing – original draft: SW, JB, CSS

Writing – review & editing: SW, JB, KA, VP, JT, TM, AL, CSS

## Data and materials availability

All data are available in the main text or the supplementary materials. Raw microscopical data are available upon request. The mass spectrometry data have been deposited and will be available upon acceptance.

## Supplemental figure legends

**Figure S1:** (A) Progression throughout the cell cycle is strictly regulated by the activity and level of cyclins and cyclin-dependent kinases (CDKs). CDK1 is essential for the transition between the G2 phase and mitosis. CDK1 can alone drive the cell cycle, such as in G1/S transition upon CDK2 loss. CDK1 is targeted by WEE1 and PKMYT1 kinases to balance CDK1 activity levels through the cell cycle. WEE1 catalyzes inhibitory phosphorylation on Tyrosine 15 of both CDK1 and CDK2, limiting kinase activities. PKMYT1 phosphorylates CDK1 at Threonine 14 to reduce CDK1 activity.

(B) Example of editing outcomes at the CDK1 T14-Y15 target site in one of the short term CRISPR-PTM analysis (Fig 1E) of CDK1-Y15 phospho-mimetic and non-phosphorylatable mutations (CDK1-Y15F, CDK1-Y15E). CDK1-T14A/Y15F (CDK1 AF) was included as positive control. Data show read values between day 1 and day 5.

(C) CRISPR-PTM analysis of different combinations of non-phosphorylatable mutations (CDK1-T14A, Y15F and T14A/Y15F). CRISPR knock-in Cassettes were delivered to iCas9-MCF10A cells, and Mut:WT′ ratios were determined on day 2 and at day 12 and normalized to day 2 value. Data are means ± S.E.M of n = 8 independent biological replicates.

(D) Dose response matrix for cell viability upon 5-day treatment with WEE1i inhibitor (adavosertib) in combination with PKMYT1i (lunresertib) in iCas9-MCF10A, *TP53*KO. Data represent mean from technical triplicate.

(E) Immunoblotting for lot CDK1 pT14, pY15 and total CDK1 using lysates from MCF10 *TP53*KO after treatment with either adavosertib or lunresertib 250 nM for 4h.

(F) CRISPR knock-in analysis of non-phosphorylatable mutation (CDK1Y15F) with PKMYT1 inhibition. CRISPR knock-in Cassettes were delivered to iCas9-MCF10A *TP53*KO cells, and Mut:WT′ ratios were determined on 12 treated with either DMSO as control or with 250 nM lunresertib and normalized to day 12 DMSO value. Means ± S.E.M of n = 3 P values are from Paired t-test with * < 0.0332, ** < 0.0021, *** <0.0002, and **** < 0.0001.

**Figure S2:**

**(A)** Validation of MCF10A *CDK2*KO by immunoblotting. Representative image of n = 3 independent biological replicates.

(B) Dose response matrix for cell viability upon 5-day treatment with WEE1i inhibitor (adavosertib) in combination with PKMYT1i (lunresertib) in iCas9-MCF10A, *TP53*KO, *CDK2* KO cells, data represent mean from technical triplicate.

(C) Immunoblotting analysis for CDK1 pT14, pY15 and total CDK1 using lysates from MCF10A *TP53*KO *CDK2* KO after treatment with either adavosertib or lunresertib 250 nM for 4h.

**Figure S3:**

**(A)** Plot of CDK1 T14, Y15 and Y19 phosphorylation status after treatments either with adavosertib (1 µM, 90 min) or lunresertib (1 µM, 90 min) followed by phosphoproteomics.

(B) Immunoblotting to detect controls of WEE1i (1 μM, 4 h) and lambda phosphatase (PPase) treatment based on the CDK1 pY15 site in iCas9-MCF10A *TP53*KO, *CDK2* KO cells. Means ± S.E.M of n=3 (WEE1i) or 4 (PPase) independent biological replicates.

(C) Immunoblotting the CDK1-Y19 phosphorylation site using CDK1 pY19 specific antibody in U2OS cells with non-treated sample (NT), WEE1i (1 μM, 4 h) and PPase treatment by Western blot. The endogenous signal for CDK1-pY19 is marked with an arrow. Means ± S.E.M of n = 3 (adavosertib) or 4 (PPase) independent biological replicates.

(D) Immunoblotting for endogenous CDK1 pY19, 48 h after siRNA CDK1 treatment. The endogenous signal for CDK1-pY19 is marked with an arrow. Means ± S.E.M; n = 3

(E) CRISPR knock-in analysis of different combinations of non-phosphorylatable mutations (CDK1-Y19F). CRISPR knock-in cassettes were delivered to iCas9-MCF10A *TP53*KO, *CDK2* WT or CDK2 KO cells. Mut:WT′ ratios were determined on 12 treated with either DMSO as control or with 250 nM lunresertib and normalized to the day 12 DMSO value. Means ± S.E.M ; n = 4

(F) CRISPR knock-in analysis of different combinations of non-phosphorylatable mutations (CDK1-T14A/Y19F). CRISPR knock-in Cassettes were delivered to iCas9-MCF10A *TP53*KO *CDK2* WT or *CDK2* KO cells, and Mut:WT′ ratios were determined on day 2 and at day 12 and normalized to day 2 value. Means ± S.E.M; n = 4

P values are from Paired t-test with * = 0.0332, ** = 0.0021, *** = 0.0002, and **** = 0.0001.

**Figure S4:**

**(A)** RCA per cell in transfected cells with CRISPR knock-in cassette or in non-transfected cells for WT’, CDK1-Y15F and CDK1-Y19F probes. The percentage represents positive cells counted. Representative experience for n = 3 independent biological replicates:

(B) Percentage of edited cells detected by CRISPR-VEIS compared to NGS result. Data are normalized to NGS results. Means ± S.E.M; n = 3

**Figure S5:**

**Table 1:** List of gRNAs

**Table 2:** List of ssODN

**Table 3:** List of primers

**Table 4:** List of LNA primers

**Table 5:** List of padlock probes

**Table 6:** List detection oligonucleotides

## Materials and Methods

### Cells and culture conditions

Immortalized human breast epithelial cells expressing doxycycline-inducible Cas9 (iCas9-MCF10A) were a gift from Roderick L. Beijersbergen, (The Netherlands Cancer Institute). iCas9-MCF10A *TP53*KO cell line was generated as previously described^27^.

MCF10A cells were cultured in Dulbecco’s modified Eagle medium/F12, HEPES (Thermo Fisher Scientific, 31330038) supplemented with 5% (vol/vol) horse serum (Thermo Fisher Scientific, 26050088), 20 ng/ml EGF (Peprotech, AF-100-15), 10 μg/ml insulin (Sigma, I1882), 0.5 μg/ml hydrocortisone (Sigma, H0888) and 100 ng/ml cholera toxin (Sigma, C8052). Human osteosarcoma cell line U2OS (ATCC) were cultured in Dulbecco’s Modified Eagle’s medium (DMEM; Gibco) supplemented with 10% fetal bovine serum (FBS; Cytiva) and 1% Penicillin-Streptomycin (10,000 U/mL; Gibco). All cells were cultured at 37°C in a 5% CO2 humidified incubator.

### Cell line generation

To generate the iCas9-MCF10A *TP53*KO *CDK2* KO cell line, iCas9-MCF10A *TP53*KO cells were transfected with a gRNA targeting the beginning of CDK2 (see Table 1). Transfected cells were single-cell sorted in 96-well plates 2 days after transfection, and the obtained clones were tested for CDK2 frameshift by Sanger sequencing (Eurofin) and western blot for protein levels. To create a stable iCas9-MCF10A *TP53*KO *CDK2* KO cell line that expresses doxycycline-inducible empty GFP, CDK1-WT-GFP or mutants in an inducible manner, HEK293T cells were transfected with 3 μg of either empty, WT or mutant pLVX-CDK1-GFP plasmid, 1 μg VSV-G (Clontech), and 1 μg PAX2 (Clontech) plasmids using JetPEI (Polyplus) to produce lentivirus according to the manufacturer’s protocol. 24 h post-transfection, the supernatant was collected to obtain lentiviral particles. The supernatant containing lentiviral particles was mixed with fresh medium containing 10 μg/ml polybrene (Sigma Aldrich H9268). This was added to the iCas9-MCF10A *TP53*KO *CDK2* KO cells. 24h after transduction, transduced cells were selected using 2 μg/ml puromycin for 10 days. Plasmids were synthesized and produced by GenScript (https://www.genscript.com/)

### CRISPR editing

CRISPR cassettes were designed as previously described^26^. Briefly, gRNA (Table 1) was selected for Cas9 with the online software Benchling (https://benchling.com). The base pairs to be mutated were located as close as possible to the PAM sequence to promote effective knock-in.

Repair templates were designed as single-stranded oligodeoxynucleotides (ssODNs) with 45 nucleotides homology arms flanking the codon to be mutated. Additionally, the synonymous WT′ mutation was placed at the same codon position as the variant of interest to promote knock-in at similar frequencies. If the mutation site was distant from the PAM region, a silent mutation was added closer to the PAM to improve editing efficiency, which is the case for some CDK1 repair templates. For the WT’ the Codon Usage Database (https://.genscript.com/tools/codon-frequency-table) was used to verify that the mutation did not produce a rarely used codon.

All sequences for repair templates can be found in Table 2.

gRNAs were used in the form of crRNA:tracrRNA duplexes purchased from Integrated DNA Technologies. Repair templates were purchased as unmodified Ultramer DNA oligonucleotides at 100 μM in IDTE, pH 8.0 from Integrated DNA Technologies. For all iCas9-MCF10A cells, Cas9 expression was induced by adding 1 μg/ml doxycycline to the culture medium 24 hours before transfection of 60–70% confluent cells. Cells were transfected with 30 nM crRNA, tracrRNA and 8 nM of repair templates (equal mix of WT’ and Mut) using Lipofectamine RNAiMAX (Thermo Fisher Scientific, 13778) under conditions specified by the manufacturer. After the transfection, an aliquot of cells was collected for the early time point Mut:WT’ analysis. The remaining cells were split into untreated conditions (DMSO) and treated with either WEE1i (adavosertib or zedoresertib) or PKMYT1i (lunresertib) conditions at 250 nM. The culture media was changed every 3 days, and cells were split before confluency was reached. Cells were harvested on a later day after transfection for the late time-point for the Mut:WT’ analysis.

### CRISPR Mut:WT′ analysis

Genomic DNA was extracted from CRISPR-PTM or CRISPR knock-in -edited cell populations using the following: GenElute Mammalian Genomic DNA Miniprep Kit (Sigma, G1N350-1KT) for CRISPR-PTM samples. For all PCRs, 100 ng of genomic DNA was used as a template.

Primer pairs for PCR amplification (Table 3) of the target site were designed to be at least 25 nucleotides away from the mutation of interest to generate PCR products of 150–350 base pairs, using the primer3 website (https://primer3.ut.ee/). Afterwards, the PCR product was used as the template for a second PCR, in which the primers contained overhangs with sample-specific barcodes, as well as adaptors for next-generation sequencing (NGS).

The amplicon library was prepared using a MiSeq Reagent Kit v2 (Illumina, MS-102-2002) and sequenced in a MiSeq instrument (Illumina, SY-410-1003), according to the manufacturer’s instructions. A library of 10 pM was sequenced with a concentration of 20% PhiX in the final library. NGS data were analyzed by the CRISPResso2 online tool^41^ using default settings (.crispresso.pinellolab.org/submission). Robust biological reproducibility in CRISPR-PTM experiments requires a minimum sequencing depth of 10,000 reads per sample and an editing efficiency of at least 1 % for the targeted mutation.

### Cell growth modelling

#### Data Processing and Scaling

Cells were transfected with CDK1 crRNA:tracrRNA duplexes together with an equal ratio mix of repair template containing for each phosphosite (T14 or Y15) a non-phosphorylatable mutant, phospho-mimetic, WT’ and CDK1 AF (For example, T14 site had: T14A, T14D, T14T and CDK1 AF). Raw experimental measurements for cell populations (wild-type and mutants) were obtained every 24h after transfection for a total of 5 days. Datapoints were normalized to estimated cell counts using a predefined mapping of days to cell numbers assuming exponential proliferation (1: 50, 2: 100, 3: 200, 4: 400, 5: 800). For each condition, scaled values were computed by multiplying the observed read counts by the corresponding cell count factor.

#### Mathematical Models

Two growth models were applied to account for a biphasic growth pattern^33^:

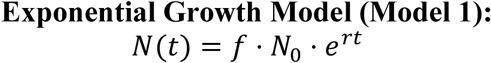

1. where 𝑁(𝑡) is the population size at time 𝑡, 𝑁_0_ is the initial population, 𝑓is the fraction of dividing cells, and 𝑟is the growth rate. 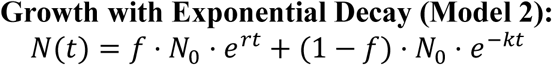
2. where 𝑘 is the decay constant accounting for non-dividing or dying cells.

#### Parameter Estimation

Nonlinear least-squares fitting was performed using the curve fit function from **SciPy**^42^ (version 1.11.4) in Python (version 3.9.19). Model 1 parameters (𝑓, 𝑟) were estimated from early time points (≤ day 2), while Model 2 parameters (𝑓, 𝑟, 𝑘) were estimated from later time points (≥ day 2). For wild-type, 𝑓was fixed at 1.0 during Model 2 fitting. Initial guesses and bounds were applied to ensure biologically plausible estimates.

#### Model Evaluation

Goodness-of-fit was assessed using the coefficient of determination (𝑅^2^) and Akaike Information Criterion (AIC)

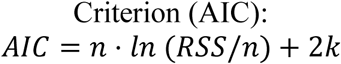

where 𝑛 is the number of observations, RSS is the residual sum of squares, and 𝑘is the number of parameters.

#### Fitness

Relative fitness was calculated based on data from Model 2 for the span from day 2 to day 5 using the formula

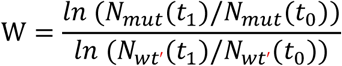

### Western Blot and GFP-immunoprecipitation

For protein extraction, cells were lysed in RIPA (Sigma) buffer containing EDTA-free protease inhibitor cocktail (Roche) and phosphatase inhibitors (Roche) for 30 min on ice. Lysates were sonicated for a total of 10 cycles, 30 s on/off. Lysates were centrifuged for 15 min at 16 000 × g at 4°C. Protein concentration was then measured with Bradford assay and adjusted to ensure equal loading.

For GFP-immunoprecipitation, 500 µg of the protein was used further for immunoprecipitation. GFP-Trap beads (ChromoTek, gta-100) were equilibrated by suspending 25 µl of bead slurry per IP reaction in 500 µl of RIPA lysis buffer. The beads were washed twice with lysis buffer. The protein lysate was added to the GFP beads and incubated at 4°C for 2 hours with constant mixing. Following incubation, the tubes were centrifuged at 2000 × g for 2 minutes at 4°C, and the supernatant was discarded. The beads were washed three times with 1 ml of lysis buffer. For Lambda phosphatase data points, GFP beads were mixed in RIPA without phosphatase inhibitor and with 1 μL of Lambda phosphatase (NEB) and phosphatase buffer at 30°C for 30min.

For other protein Lysates or GFP beads, samples were mixed with 4× Laemmli sample buffer (Sigma) complemented with NuPAGE reducing agent and boiled for 10 min at 95°C. Samples were run on NuPAGE Bis-Tris 4-12% gels according to manufacturer instructions. Proteins were then transferred to a nitrocellulose membrane and blocked with PBS + 0.1% Tween 20 + 5% Milk powder (Sigma) and incubated overnight with primary antibodies at 4°C. The membrane was then washed 3 × 5 min in PBS + 0.1% Tween20 and incubated with secondary HRP-conjugated antibodies (Vector Laboratories) for 1 at room temperature. Membranes were washed 3 × 5 min with PBS + 0.1% Tween20 and incubated with Classico Western HRP substrate (Millipore-Sigma) for 2 min. Chemiluminescence signal was detected using a Bio-Rad ChemiDoc Touch Imaging System. Images were analyzed and quantified using Licor Image Studio software.

The following primary antibodies were used

Vinculin 1:10000, Sigma Cat#V9131, CDK1 1:1000, Abcam Cat#ab18; CDK1pY15 1:1000, Cell Signaling Technology, Cat#9111S, CDK1pT14 1:1000, Abcam, Cat#ab58509, GFP 1:20000, Abcam AB-18.0107, CDK1 pY19, 1:500 custom-made (this study) by Moravian Biotechnology.

### Mass spectrometry

Mass Spectrometry analysis was performed as described previously^43^. Briefly, U2OS cells were treated for 90 minutes (n = 3) with DMSO, 1 µM of adavosertib, 1 µM of lunresertib or 1µM combination of the two. Afterwards, the cells were subsequently harvested by scraping. Harvested cells were lysed using 100 µl of lysis buffer (6 M Guanidinium Hydrochloride, 10 mM TCEP, 40 mM CAA, 50 mM HEPES pH 8.5). Samples were boiled at 95°C for 5 minutes, after which they were sonicated on high for 5× 30 seconds in a Bioruptor sonication water bath (Diagenode) at 4°C. After determining protein concentration with Pierce™ Rapid Gold BCA Protein Assay Kit (Thermo Fisher), 200 ug were taken forward for digestion. Samples were diluted 1:3 with 10% Acetonitrile, 50 mM HEPES pH 8.5, LysC (MS grade, Wako) was added in a 1:50 (enzyme to protein) ratio, and samples were incubated at 37°C for 4 hours. Samples were further diluted to 1:10 with 10% Acetonitrile, 50 mM HEPES pH 8.5, trypsin (MS grade, Promega) was added in a 1:100 (enzyme to protein) ratio, and samples were incubated overnight at 37°C. Enzyme activity was quenched by adding 2% trifluoroacetic acid (TFA) to a final concentration of 1%. Prior to TMT labeling, the peptides were desalted with the use of SOLAµ™ SPE Plate (Thermo Fisher). The sorbent was activated by 40 µl of 100% Methanol (HPLC grade, Sigma), then 40 µl of 80% Acetonitrile, 0.1% formic acid. The plate was subsequently equilibrated 2 × with 40 µl of 1% TFA, 3% Acetonitrile, after which the samples were loaded. The plates were then washed 2 times with 200 µL of 0.1% formic acid, and the bound peptides were eluted into clean 500 µl Eppendorf tubes using 40% Acetonitrile, 0.1% formic acid. The eluted peptides were concentrated in an Eppendorf Speedvac and reconstituted in 50mM HEPES (pH 8.5).

Ti-IMAC HP (ReSyn Bioscience) was used for enrichment of phosphopeptides, 50 µL of the particles were equilibrated with 200 µL of 70 % ethanol for 5 min and subsequently with 200µL 1 % NaNH3 solution for 10 min. Particles were then washed three times with 100 µL of 80% Acetonitrile, 1M glycolic acid, 5% TFA.

Resuspended peptides were diluted 1:1 with 80% Acetonitrile, 1M glycolic acid, 5% TFA were loaded onto the particles and incubated for 30 min with gentle agitation. Supernatant was then removed and kept as the proteome sample. The particles were washed twice with 400 µL of 80% Acetonitrile, 1% TFA for 2 min and twice with 400 µL of 10% Acetonitrile 0.1% TFA. The phosphopeptides were then eluted in three rounds by adding 100 µL of 1 % NaNH3 solution for 20min. The eluted phosphopeptide solution was then acidified with 40 µL of TFA, and the supernatant proteome samples were diluted to reduce the Acetonitrile concentration to 5%. Peptide solutions were then desalted with the use of SOLAµ™ SPE Plate, and subsequently peptides were concentrated in an Eppendorf Speedvac. Concentrated peptides were stored at -80°C. The desalted peptide samples were resuspended in 2% CAN, 0.1% TFA and 500 ng were loaded onto EvoSep stagetips according to manufacturer’s protocol.

Both phosphoproteome and proteome peptides were analyzed with a “20 samples per day” method on the EvoSep One instrument and analyzed on Q-Exactive Exploris 480 instrument (Thermo Fisher Scientific) running a high-resolution MS1 (HRMS1) data-independent acquisition method. Full MS spectra were collected at 120 000 with AGC target of 3 × 10^6 or maximum injection set to auto. Scan range from 400-1000 m/z was used. The MS2 spectra were obtained with 60,000 resolution, with an AGC target of 10e5. 75 windows of 8 m/z, with 1m/z overlap, were used with a MS1 scan every 200 m/z. FAIMS CV was set -45. The raw files were the analysis with Spectronaut 16 software.

### Cell Viability assay

Drugs were dispensed into 96-well plates (Greiner-BIO), and cells were dispensed at a desired number of 100 μL. After 5 days of incubation at 37°C, 30 μL of PBS containing 4 μg/mL Hoechst 33342 and 1/10 000 CellTox Green dyes were added for 1 hour prior to imaging. Images were obtained automatically with the ScanR acquisition software using Olympus Universal Plan Super Apo 4 × / 0.16 AIR Objective.

### CRISPR-VEIS

#### Design of the probes and primers

Reverse transcription LNA primer (Table 4), Padlock probes and detection nucleotides were designed as described previously^38^ and specific sequences can be found in the following table. All oligonucleotides were ordered with IDT.

To ensure specificity of the padlock probe between single-nucleotide variants, the probe is designed in a way that the mutation of interest is placed at the 3′ end. Only probes with a perfect match on the 3’ end will be ligated into a circular molecule. For CDK1 MUT and WT’ probes, several silent mutations were introduced with the repair template, ensuring high specificity of the probe. Probes were ordered pre-phosphorylated from IDT (Table 5).

After rolling circle amplification, the probe was detected using detector probes conjugated with Alexa fluor. Detector probes were ordered with IDT (Table 6).

#### CRISPR-VEIS mRNA staining

5 Day after CRISPR knock-in transfection, cells were seeded on coverslips previously coated with Poly-L-Lysine Solution (Sigma). After 24 hours, cells were fixed in 4% paraformaldehyde for 10-20 min and washed twice in PBS-Tween 0.05 (PBS-T) before a permeabilization step with 0.25% Triton X-100 for 5 min. Cells were washed twice in PBS-T before proceeding with the Reverse Transcription. Reverse transcription, Padlock probe ligation, rolling circle amplification (RCA) and detection were done as previously described^28^. Briefly, Reverse transcription was done at 45°C for 1 h using 5 U/µL RevertAid minus M-MuLV reverse transcriptase (Thermo), 5 U/µL RiboLock RNase inhibitor (Thermo), 1 × M-MuLV RT buffer, 0.5 nm dNTP (NEB), 0.2 µg/µL BSA (NEB) and 50 nM LNA primer for CDK1 (IDT). Then coverslips were washed twice with PBS-T before being fixed for a second time with PFA 4% for 10 min at RT followed by 2 PBS-T washes.

RNAse digestion and probe hybridization were done for 30 min at 37°C, followed by 30 min at 45°C using 0.5 U/µL of Ampligase (Biosearch Technologies), 0.1 U/µL of RNaseH (NEB), 1 × Ampligase buffer, 2 µg/µL BSA (NEB), 0.05M KCl, 20% formamide and 0.1 μM padlock probe. Rolling circle amplification was done with 2.5 U/µL of Phi29 DNA polymerase (NEB), 1 × Phi29 buffer, 0.25nm dNTP (NEB), 2 U/µL BSA (NEB) and 5% glycerol. Coverslips were incubated for 1 h at 37°C.

Detection was done with 0.1 μM of Detection oligonucleotide coupled with either Alexa 488 or 594. All steps were done on coverslips in a sealed humid chamber with 30 µL working volume. Cells were then immunolabeled using FOXM1pT600 1:1000, (Cell Signaling) antibodies diluted with PBS-T-1% BSA for 1 hour at RT. Cells were then probed with a secondary anti-rabbit antibody coupled to Alexa Fluor 647 (Life Technologies), and DNA was stained with DAPI 1 µg.mL^-1^ (Sigma-Aldrich) and mounted with fluorescent mounting medium (Fluoromount-G, Southern Biotech).

### Quantitative image-based cytometry

Images used for QIBC were obtained with the ScanR acquisition software controlling a motorized Olympus IX-83 wide-field microscope. The system was equipped with filter cubes compatible with DAPI, FITC, Cy3, and Cy5 fluorescent dyes. Unless stated otherwise, the QIBC data presented in this study were obtained using an Olympus Universal Plan Super Apo 20 × Objective. The acquired images were processed using the ScanR image analysis software.

For CRISPR-VEIS, cells with at least 2 foci in the cytoplasm were considered as positive for the mutation of interest. A minimum of 150 positive cells were used for quantification per replicate.

### Statistical analysis and figure design

We used a paired t-test to calculate the significance in all cases. We used GraphPad Prism 10 (GraphPad Prism version 10 for Windows, GraphPad Software, graphpad.com) to generate the data plots and do the statistical analysis. Data distribution was assumed Gaussian, but this was not tested. Explanatory panel figures were made using Biorender (biorender.com), PowerPoint or Adobe Illustrator.

