## Supplemental figures for "CRISPR-PTM and CRISPR-VEIS: Multiplexed platforms for quantitative functional analysis of endogenous phosphosites"

Supplemental Figure 1

A

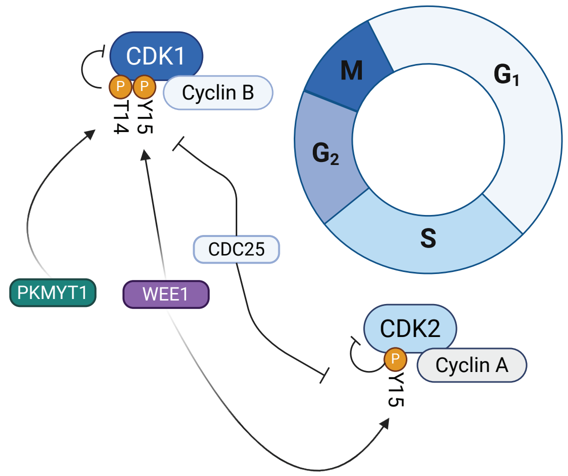


B

| | WT/Unmodified | CDK1 Y15F | CDK1 Y15E | CDK1 AF | WT' | Frameshift inDels | Inframe mutation | Total Mutations | CDK1 Y15F:WT' ratio | CDK1 Y15E:WT' ratio | CDK1 AF:WT' ratio | Total reads |
|---|---|---|---|---|---|---|---|---|---|---|---|---|
| Day 1 | 43.7% | 2.23% | 10% | 1.87% | 2.99% | 47.1% | 9.2% | 56.3% | 74.40% | 318.50% | 62.38% | |
| Reads | 13964 | 712 | 3048 | 597 | 957 | 15069 | 2929 | 17998 | | | | 31962 |
| Day 5 | 85.1% | 4.57% | 3.18% | 0.09% | 5.86% | 10% | 5% | 15% | 74.79% | 34.54% | 1.52% | |
| Reads | 14640 | 786 | 900 | 16 | 1658 | 1717 | 849 | 2566 | | | | 17206 |

C

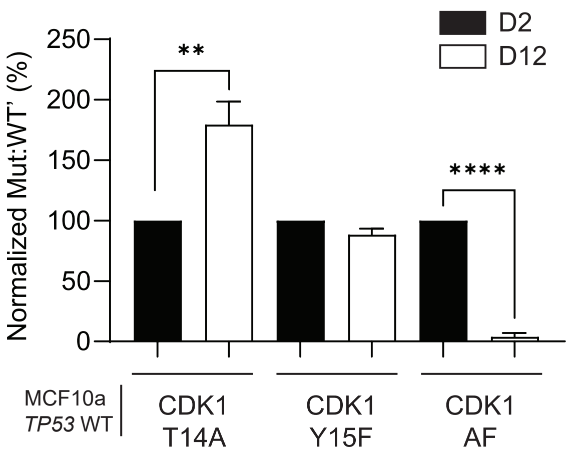


D

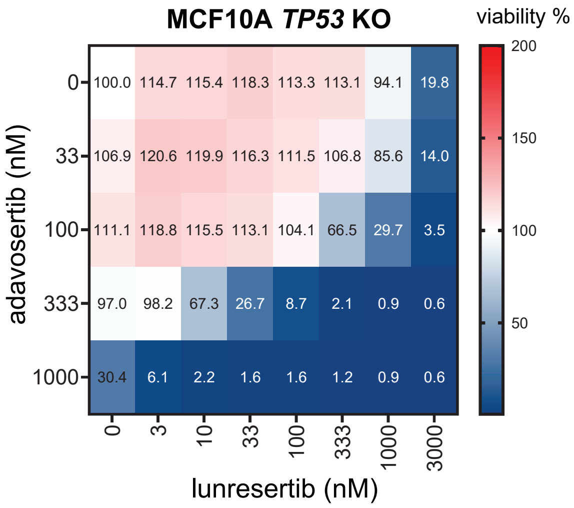


E

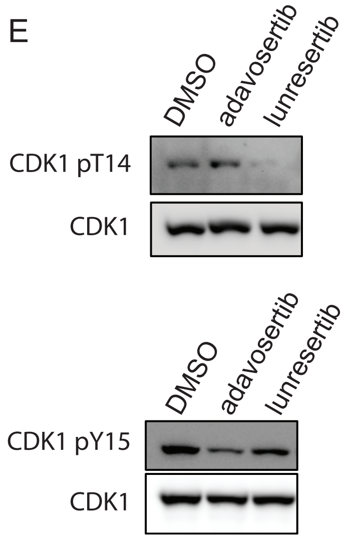


F

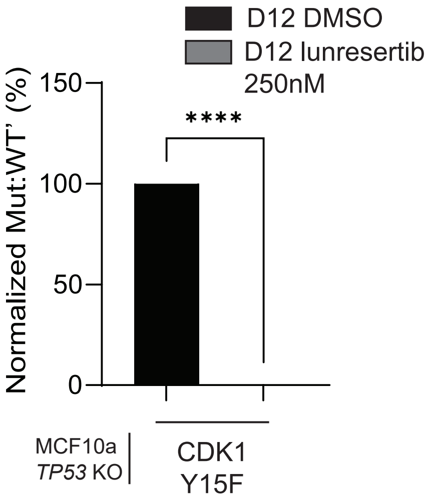

Supplemental Figure 2

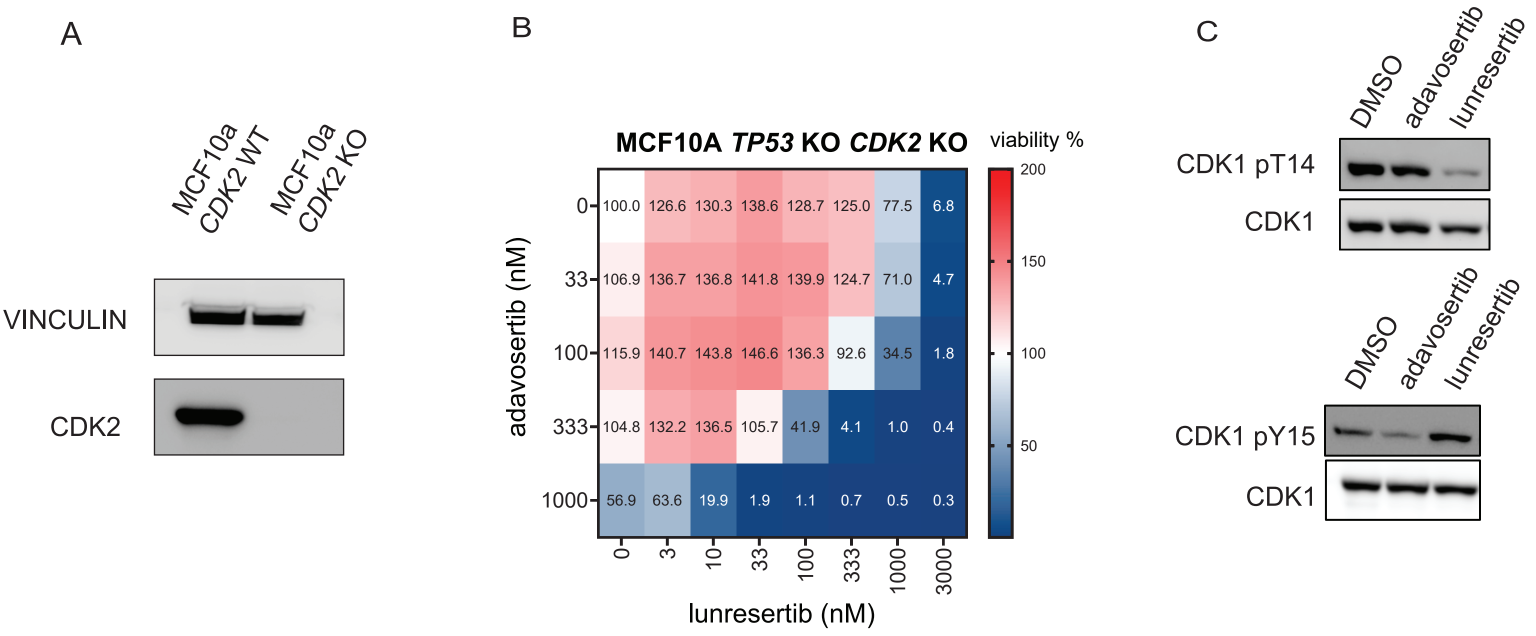

Supplemental Figure 3

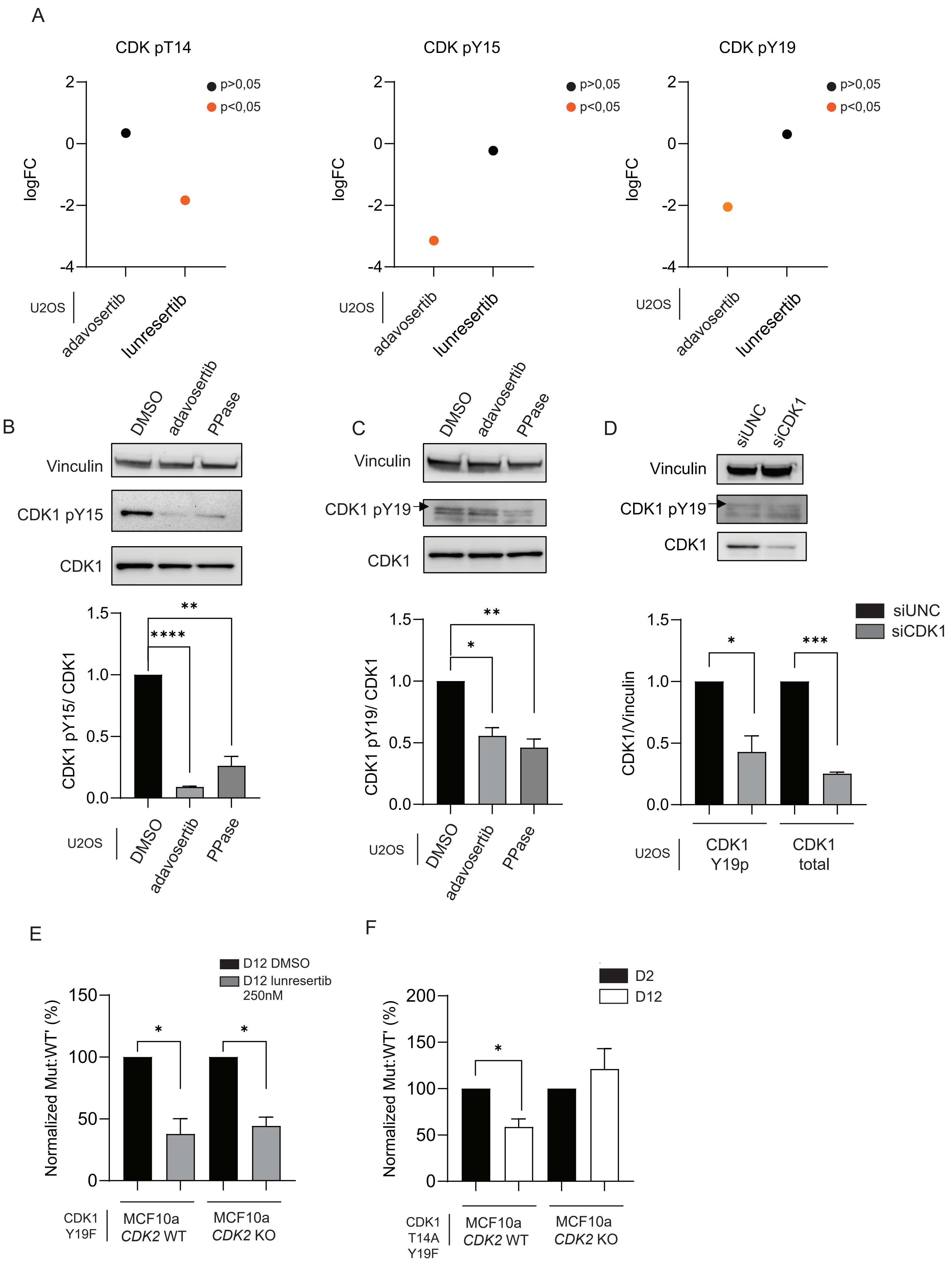

Supplemental Figure 4

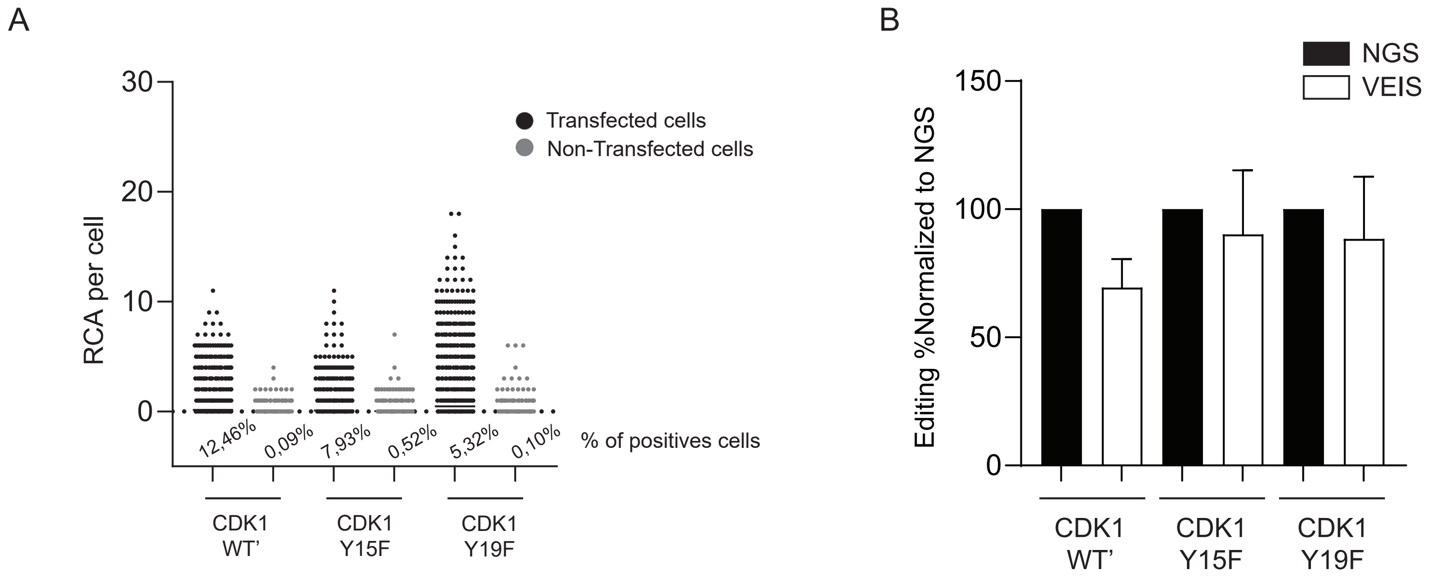

Table 1

Supplemental Figure 5

| Gene | gRNA sequence (5'-3') |
|---|---|
| CDK1 | TTTACTTTGTTTCAGGTACCTA |
| CDK2 | AAGATCGGAGAGGGCACGTA |

Table 2

| | | Repair template sequence |
|---|---|---|
| CDK2 AF | Mut | TTCATGGAGAACTTCCAAAAGGTGGAAAAGATCGGAGAGGGCGCGTTCGGAGTTGTGTACAAAGCCAGAAACAAGTTGACGGGAGAGGTGGTG |
| | WT' | TTCATGGAGAACTTCCAAAAGGTGGAAAAGATCGGAGAGGGAACCTACGGAGTTGTGTACAAAGCCAGAAACAAGTTGACGGGAGAGGTGGTG |
| CDK1 AF | Mut | GTGGGGTGTGTCACACAGCATATTATTTACTTTGTTTCAGGTGCCTTTGGAGTCGTGTATAAGGGTAGACACAAAACTACAGGTCAAGTGGTA |
| | WT' | GTGGGGTGTGTCACACAGCATATTATTTACTTTGTTTCAGGTACGTACGGAGTCGTGTATAAGGGTAGACACAAAACTACAGGTCAAGTGGTA |
| CDK1 T14A | Mut | GTGGGGTGTGTCACACAGCATATTATTTACTTTGTTTCAGGTGCCTATGGAGTCGTGTATAAGGGTAGACACAAAACTACAGGTCAAGTGGTA |
| | WT' | GTGGGGTGTGTCACACAGCATATTATTTACTTTGTTTCAGGTACGTATGGAGTCGTGTATAAGGGTAGACACAAAACTACAGGTCAAGTGGTA |
| CDK1 Y15F | Mut | GTGGGGTGTGTCACACAGCATATTATTTACTTTGTTTCAGGTACCTTTGGAGTCGTGTATAAGGGTAGACACAAAACTACAGGTCAAGTGGTA |
| | WT' | GTGGGGTGTGTCACACAGCATATTATTTACTTTGTTTCAGGTACCTACGGAGTCGTGTATAAGGGTAGACACAAAACTACAGGTCAAGTGGTA |
| CDK1 Y19F | Mut | GTGGGGTGTGTCACACAGCATATTATTTACTTTGTTTCAGGTACCTATGGAGTTGTGTTTAAGGGTAGACACAAAACTACAGGTCAAGTGGTA |
| | WT' | GTGGGGTGTGTCACACAGCATATTATTTACTTTGTTTCAGGTACCTATGGAGTTGTGTACAAGGGTAGACACAAAACTACAGGTCAAGTGGTA |
| CDK1 Y15F-Y19F | Mut | GTGGGGTGTGTCACACAGCATATTATTTACTTTGTTTCAGGTACCTTTGGAGTTGTGTTTAAGGGTAGACACAAAACTACAGGTCAAGTGGTA |
| | WT' | GTGGGGTGTGTCACACAGCATATTATTTACTTTGTTTCAGGTACCTACGGAGTTGTGTACAAGGGTAGACACAAAACTACAGGTCAAGTGGTA |
| CDK1 T14A-Y19F | Mut | GTGGGGTGTGTCACACAGCATATTATTTACTTTGTTTCAGGTGCCTATGGAGTTGTGTTTAAGGGTAGACACAAAACTACAGGTCAAGTGGTA |
| | WT' | GTGGGGTGTGTCACACAGCATATTATTTACTTTGTTTCAGGTACGTATGGAGTTGTGTACAAGGGTAGACACAAAACTACAGGTCAAGTGGTA |
| CDK1 DE | Mut | GTGGGGTGTGTCACACAGCATATTATTTACTTTGTTTCAGGTGACGAAGGAGTCGTGTATAAGGGTAGACACAAAACTACAGGTCAAGTGGTA |
| | WT' | GTGGGGTGTGTCACACAGCATATTATTTACTTTGTTTCAGGTACGTACGGAGTCGTGTATAAGGGTAGACACAAAACTACAGGTCAAGTGGTA |
| CDK1 T14D | Mut | GTGGGGTGTGTCACACAGCATATTATTTACTTTGTTTCAGGTACCTATGGAGTTGTGTATAAGGGTAGACACAAAACTACAGGTCAAGTGGTA |
| | WT' | GTGGGGTGTGTCACACAGCATATTATTTACTTTGTTTCAGGTACCTATGGAGTTGTGTATAAGGGTAGACACAAAACTACAGGTCAAGTGGTA |
| CDK1 Y15E | Mut | GTGGGGTGTGTCACACAGCATATTATTTACTTTGTTTCAGGTACCGAAGGAGTCGTGTATAAGGGTAGACACAAAACTACAGGTCAAGTGGTA |
| | WT' | GTGGGGTGTGTCACACAGCATATTATTTACTTTGTTTCAGGTACCTACGGAGTCGTGTATAAGGGTAGACACAAAACTACAGGTCAAGTGGTA |
| CDK1 Y19E | Mut | GTGGGGTGTGTCACACAGCATATTATTTACTTTGTTTCAGGTACCTATGGAGTTGTGGAAAAGGGTAGACACAAAACTACAGGTCAAGTGGTA |
| | WT' | GTGGGGTGTGTCACACAGCATATTATTTACTTTGTTTCAGGTACCTATGGAGTTGTGTACAAGGGTAGACACAAAACTACAGGTCAAGTGGTA |
| CDK1 Y15E-Y19E | Mut | GTGGGGTGTGTCACACAGCATATTATTTACTTTGTTTCAGGTACCGAAGGAGTTGTGGAAAAGGGTAGACACAAAACTACAGGTCAAGTGGTA |
| | WT' | GTGGGGTGTGTCACACAGCATATTATTTACTTTGTTTCAGGTACCTACGGAGTTGTGTACAAGGGTAGACACAAAACTACAGGTCAAGTGGTA |
| CDK1 T14D-Y19E | Mut | GTGGGGTGTGTCACACAGCATATTATTTACTTTGTTTCAGGTGACTATGGAGTTGTGGAAAAGGGTAGACACAAAACTACAGGTCAAGTGGTA |
| | WT' | GTGGGGTGTGTCACACAGCATATTATTTACTTTGTTTCAGGTACGTATGGAGTTGTGTACAAGGGTAGACACAAAACTACAGGTCAAGTGGTA |

Table 3

| Primer name | Primer Sequence (5'-3') |
|---|---|
| CDK1 FWD | 5’- ACACTCTTTCCCTACACGACGCTCTTCCGATCTCTTTAGTTTGTGGGGTGTGTCA -3’ |
| CDK1 REV | 5’- TGACTGGAGTTCAGACGTGTGCTCTTCCGATCTTCCCGAATTGCAGTACTAGGA -3’ |
| CDK2 FWD | 5’- ACACTCTTTCCCTACACGACGCTCTTCCGATCTGCTGGCGCTTCATGGAGA -3’ |
| CDK2 REV | 5’- TGACTGGAGTTCAGACGTGTGCTCTTCCGATCTGAAGTCCCTCCCTGCTCTC -3’ |

Table 4

| RT primer | Primer Sequence (5'-3') |
|---|---|
| CDK1 | C+CA+CT+TG+AC+CT+GT+AGTTTTGTGT |

Table 5

| Padlock probes | Padlock probes Sequence (5'-3') |
|---|---|
| Padlock CDK1-WT | PHOSPHO____TAAGGGTAGACACAATTCCTTTTACGACCTCAATGCACATGTTTGGCTCCTCTT CCTATGGAGTTGTGTA |
| Padlock CDK1-Y15F | PHOSPHO____ GTGTATAAGGGTAGATTCCTTTTACGACCTCAATGCACATGTTTGGCTCCTCTT AGGTACCTTTGGAGTC |
| Padlock CDK1-Y19F | PHOSPHO____TAAGGGTAGACACAAATTCCTTTTACGACCTCAATGCACATGTTTGGCTCCTCTT CGTACGGAGTCGTGTT |
| Padlock CDK1-WT' | PHOSPHO____GTGTATAAGGGTAGATTCCTTTTACGAAGTAGCCGTGACTATCGACTTCTTAGGTACGTACGGAGTC |

Table 6

| | Detection oligonucleotide (5'-3') |
|---|---|
| Decorator 1 (Alexa 488) | CCTCAATGCACATGTTTGGCTCC |
| Decorator 2 (Alexa 594) | AGTAGCCGTGACTATCGACT |
